# Characterisation of *Trichuris muris* secreted proteins and extracellular vesicles provides new insights into host-parasite communication

**DOI:** 10.1101/128629

**Authors:** Ramon M. Eichenberger, Md Hasanuzzaman Talukder, Matthew A. Field, Phurpa Wangchuk, Paul Giacomin, Alex Loukas, Javier Sotillo

**Author notes:** Both authors equally contributed to the manuscript (Ramon M. Eichenberger, Md Hasanuzzaman Talukder). Corresponding Authors: Javier Sotillo. Centre for Biodiscovery and Molecular Development of Therapeutics, James Cook University, Cairns, 4878, Queensland, Australia. Prof. Alex Loukas. Centre for Biodiscovery and Molecular Development of Therapeutics, James Cook University, Cairns, 4878, Queensland, Australia.

## Abstract

Whipworms are parasitic nematodes that live in the gut of more than 500 million people worldwide. Due to the difficulty in obtaining parasite material, the mouse whipworm *Trichuris muris* has been extensively used as a model to study human whipworm infections. These nematodes secrete a multitude of compounds that interact with host tissues where they orchestrate a parasitic existence. Herein we provide the first comprehensive characterisation of the excretory/secretory products of *T. muris.* We identify 148 proteins secreted by *T. muris* and show for the first time that the mouse whipworm secretes exosome-like extracellular vesicles (EVs) that can interact with host cells. We use an Optiprep^®^ gradient to purify the EVs, highlighting the suitability of this method for purifying EVs secreted by a parasitic nematode. We also characterise the proteomic and genomic content of the EVs, identifying >350 proteins, 56 miRNAs (22 novel) and 475 full-length mRNA transcripts mapping to *T. muris* gene models. Many of the miRNAs putatively mapped to mouse genes involved in regulation of inflammation, implying a role in parasite-driven immunomodulation. In addition, for the first time to our knowledge, we use colonic organoids to demonstrate the internalisation of parasite EVs by host cells. Understanding how parasites interact with their host is crucial to develop new control measures. This first characterisation of the proteins and EVs secreted by *T. muris* provides important information on whipworm-host communication and forms the basis for future studies.

## 1. Introduction

Infections with soil-transmitted helminths (STH) affect more than 1.5 billion people worldwide, causing great socio-economic impact as well as physical and intellectual retardation [1]. Among the STH, hookworms (*Necator americanus* and *Ancylostoma duodenale*), roundworms (*Ascaris lumbricoides*) and whipworms (*Trichuris trichiura*) are of particular importance due to their high prevalence and disease burden in impoverished countries [2]. For instance, *T. trichiura* alone infects around 500 million people worldwide, and contributes to 638,000 years of life lived with disability (YLDs) [2].

Infection with *Trichuris* spp. occurs after ingestion of infective eggs, which hatch in the caecum of the host. Larvae penetrate the mucosal tissue where they moult to become adult worms and reside for the rest of their lives. Due to the difficulty in obtaining parasite material to study whipworm infections, particularly adult worms, the rodent whipworm, *Trichuris muris,* has been extensively used as a tractable model of human trichuriasis [3, 4, 5]. In addition to parasitologists, immunologists have also benefited from the study of *T. muris* infections, and a significant amount of basic immunology research has been conducted using this model (reviewed by [6]). For instance, the role of IL-13 in resistance to nematode infections was elucidated using *T. muris* [7].

The recent publication of the genome and transcriptome of *T. muris* has provided meaningful insights into the immunobiology of whipworm infections [8]. This work provided new information on potential drug targets against trichuriasis and elucidated important traits that drive chronicity. Despite this progress, and the tractability of the *T. muris* model, very few proteomic studies have been conducted, and only a handful of reports have described proteins secreted by *Trichuris spp.* [9, 10, 11, 12, 13, 14]. Drake *et al.* characterised a pore-forming protein that *T. muris* [14] and *T. trichiura* [13] use to drill holes in the host cell membrane. Furthermore, it has been suggested that a thioredoxin-like protein secreted by the pig whipworm *Trichuris suis* plays a role in mucosal homeostasis [11].

The importance of excretory/secretory (ES) products in governing host parasite interactions and ensuring parasite survival in inhospitable environments is indisputable. Traditionally, ES products were believed to contain only soluble proteins, lipids, carbohydrates and genomic content; however, the recent discovery of extracellular vesicles (EVs) secreted by helminths has revealed a new paradigm in the study of host-parasite relationships [15, 16, 17]. Helminth EVs have immunomodulatory effects and contribute to pathogenesis. For instance, EVs secreted by parasitic flatworms can promote tumorigenesis [18] and polarise host macrophages towards a M1 phenotype [19], while EVs from the gastrointestinal nematode *Heligmosomoides polygyrus* contain small RNAs that can modulate host innate immunity [20].

In the present study, we aim to characterise the factors involved in *T. muris* host relationships. We provide the first proteomic analysis of the soluble proteins present in the ES products and we describe the proteomic and nucleic acid content of EVs secreted by whipworms. This work provides important information on whipworm biology and contributes to the development of new strategies and targets to combat nematode infections in humans and animals.

## 2. Experimental procedures

### 2.1 Ethics statement

The study was approved by the James Cook University (JCU) Animal Ethics Committee (A2213). Mice were maintained at the JCU animal house (Cairns campus) under normal conditions of regulated temperature (22°C) and lighting (12 h light/dark cycle) with free access to pelleted food and water. The mice were kept in cages in compliance with the Australian Code of Practice for the Care and Use of Animals for Scientific Purposes.

### 2.2 Parasite material, isolation of ES products, and EV purification

Parasites were obtained from genetically susceptible B10.BR mice infected with 200 *T. muris* eggs. The infection load (200 *T. muris* eggs per mouse) is well tolerated, and results, usually, in ~180 adult parasites/mouse (90% success). Adult worms were harvested from the caecum of infected mice 5 weeks after infection, washed in PBS containing 5× antibiotic/antimycotic (AA) and cultured in 6-well plates for 5 days in RPMI containing 1× AA, at 37°C and 5% CO_2_. Each well contained ~500 worms in 4.5 mL media. The media obtained during the first 4 h after parasite culturing was discarded for further analysis. Dead worms were removed and ES products were collected daily, subjected to sequential differential centrifugation at 500 *g,* 2,000 *g* and 4,000 *g* for 30 min each to remove eggs and parasite debris. For the isolation of ES products, media was concentrated using a 10 kDa spin concentrator (Merck Millipore, USA) and stored at 1.0 mg/ml in PBS at -80°C until required.

For the isolation of EVs, the media obtained after differential centrifugation was processed as described previously [21]. Briefly, ES products were concentrated using a 10 kDa spin concentrator, followed by centrifugation for 45 min at 12,000 *g* to remove larger vesicles. A MLS-50 rotor (Beckman Coulter, USA) was used to ultracentrifuge the supernatant for 3 h at 120,000 *g* and the resultant pellet was resuspended in 70 μl of PBS and subjected to Optiprep^®^ discontinous gradient (ODG) separation. One mL of 40%, 20%, 10% and 5% iodixanol solutions prepared in 0.25 M sucrose, 10 mM Tris-HCl, pH 7.2, was layered in decreasing density in an ultracentrifuge tube, and the 70 μl containing the resuspended EVs was added to the top layer and ultracentrifuged at 120,000 *g* for 18 h at 4°C. Seventy (70) μl of PBS was added to the control tube prepared as described above. A total of 12 fractions were recovered from the ODG, and the excess Optiprep^®^ solution was removed by buffer exchanging with 8 ml of PBS containing 1× EDTA-free protease inhibitor cocktail (Santa Cruz, USA) using a 10 kDa spin concentrator. The absorbance (340 nm) was measured in each of the fractions and density was calculated using a standard curve with known standards. The protein concentration of all fractions was measured using a Pierce BCA Protein Assay Kit (ThermoFischer, USA). All fractions were kept at -80°C until use.

### 2.3 Size and concentration analysis of EVs.

The size distribution and particle concentration of fractions recovered after ODG were measured using tunable resistive pulse sensing (TRPS) by qNano (Izon, USA) following the manufacturer’s instructions. Voltage and pressure values were set to optimize the signal to ensure high sensitivity. A nanopore NP100 was used for all fractions analysed except for fraction 9, where a NP150 was used. Calibration was performed using CP100 carboxylated polystyrene calibration particles (Izon) at a 1:1000 dilution. Samples were diluted 1:5 and applied to the nanopore. The size and concentration of particles were determined using the software provided by Izon (version 3.2).

### 2.4 Exosome uptake in murine colonic organoids (mini-guts)

Murine colonic organoids were produced from intestinal crypts of a female C57 Bl6/J mouse according to previous reports [22] with some modifications. Briefly, murine colonic crypts were dissociated with Gentle Cell Dissociation reagent (Stemcell Technology Inc., Canada) and further incubated in trypsin (Gibco, ThermoFischer). Approximately 500 crypts were seeded in 50 μl of Matrigel (Corning, USA) in a 24-well plate and cultured in Intesticult Organoid Growth Medium (Stemcell Technology Inc.) supplemented with 100 ng/ml murine recombinant Wnt3a (Peprotech, USA). ROCK-inhibitor (10 μM Y-27632; Sigma-Aldrich, USA) was included in the culture medium for the first 2 days to avoid anoikis.

For imaging, organoids were seeded in 75 μl of Matrigel in 6-well plates and cultured for 7 days. To investigate internalization of EVs in the colonic epithelium layer, EVs were labelled with PKH26 (Sigma-Aldrich) according to the manufacturer's instructions. A total of 15-30 million stained particles (based on the TRPS results) in 3-5 μl were injected into the central lumen of individual organoids and cultured for 3 hours at 37°C and 4°C, respectively. Cell culture medium was removed, and wells were washed with PBS. Organoids were fixed by directly adding 4% paraformaldehyde to the 6-well plates and incubating for 30 min at room temperature (RT). Matrigel was then mechanically disrupted, and cells were transferred into BSA-coated tubes. Autofluorescence was quenched by incubating the organoids with 50 mM NH_4_Cl in PBS (for 30 min at RT) and 100 mM glycine in PBS (for 5 min). Cell nuclei were stained with Hoechst dye (Invitrogen, US) and visualized on an AxioImager M1 ApoTome fluorescence microscope (Zeiss, Germany). Fluorescence intensity of PKH26-stained parasite EVs was quantified in ImageJ and expressed as percentage of corrected total fluorescence (% CTF) adjusted by background fluorescence and the surveyed area in total epithelial cells (donut-shaped selection) or in the lumen incubated at different conditions in 10 different murine colonic organoids from 2 technical replicates (5 each). The whole experiment was repeated to perform laser scanning confocal imaging on a 780 NLO microscope (Zeiss). Confocal image deconvolution was performed in ImageJ using the plugins “Diffraction PSF 3D” for PSF calculation and “DeconvolutionLab” with the Richardson-Lucy algorithm for 3D deconvolution and Tikhonov-Miller algorithm for 2D deconvolution [23].

### 2.5 Proteomic analyses

The protein content from the *T. muris* ES products and ODG fractions were analysed as follows.

#### 2.5.1 Proteomic analysis of ES products

One hundred micrograms (100 μg) of *T. muris* ES proteins from two different batches of adult worms were precipitated at -20°C overnight in ice-cold methanol. Proteins were resuspended in 50 mM NH_4_HCO_3_, reduced in 20 mM dithiothreitol (DTT, Sigma-Aldrich) and finally alkylated in 55 mM iodoacetamide (IAM, Sigma-Aldrich). Proteins were finally digested with 2 μg of trypsin (Sigma-Aldrich) by incubating for 16 h at 37°C with gentle agitation. Reaction was stopped with 5% formic acid and the sample was desalted using ZipTip^®^ (Merck Millipore). Both samples were kept at -80°C until use.

#### 2.5.2 Proteomic analysis of EVs

For the proteomic analysis of EVs, two replicates were analysed independently. The ODG fractions with a density of 1.07-1.09g/ml (fractions 5-7) were combined, and a total of 50 μg of protein from each of the two replicates was loaded on a 12% SDS-PAGE and electrophoresed at 100V for 1.5 h. Each lane was sliced into 9 pieces, which were subjected to trypsin digestion as described previously [24]. Briefly, each slice was washed for 5 min three times in 50% acetonitrile, 25 mM NH4CO3 and then dried under a vacuum centrifuge. Reduction was carried out in 20 mM DTT for 1 h at 65 °C, after which the supernatant was removed. Samples were then alkylated in 55 mM IAM at RT in darkness for 40 min. Gel slices were then washed 3× in 25 mM NH_4_CO_3_ before drying in a vacuum centrifuge followed by digestion with 500 ng of trypsin overnight at 37°C. The digest supernatant was removed from the gel slices, and residual peptides were removed from the gel slices by washing three times with 0.1% TFA for 45 min at 37°C. Samples were desalted and concentrated using Zip-Tip^®^ and kept at -80°C until use.

#### 2.5.3. Mass spectrometry and database searches

For all analyses, samples were reconstituted in 10 μl of 5% formic acid. Six microlitres of sample was injected onto a 50 mm 300 μm C18 trap column (Agilent Technologies, USA) and desalted for 5 min at 30 μL/min using 0.1% formic acid (aq). Peptides were then eluted onto an analytical nano HPLC column (150 mm x 75 μm 300SBC18, 3.5 μm, Agilent Technologies) at a flow rate of 300 nL/min and separated using a 35 min gradient (for ES proteins) or 95 min gradient (for EV proteins) of 1-40% buffer B (90/10 acetonitrile/ 0.1% formic acid) followed by a steeper gradient from 40-80% buffer B in 5 min. The mass spectrometer operated in information-dependent acquisition mode (IDA), in which a 1-s TOF MS scan from 350-1400 m/z was performed, and for product ion ms/ms 80-1400 m/z ions observed in the TOF-MS scan exceeding a threshold of 100 counts and a charge state of +2 to +5 were set to trigger the acquisition of product ion. Analyst 1.6.1 (ABSCIEX) software was used for data acquisition and analysis.

For the analysis of the ES products, a database was built using the *T. muris* genome [8] with the common repository of adventitious proteins (cRAP, http://www.thegpm.org/crap/) appended to it. A similar database containing the *T. muris* genome, the cRAP and the *Mus musculus* genome was used for the analysis of the EV mass spectrometry data Database search was performed using X!Tandem, MS-GF+, OMSSA and Tide search engines using SearchGUI [25]. Parameters were set as follows: tryptic specificity allowing two missed cleavages, MS tolerance of 50 ppm and 0.2 Da tolerance for MS/MS ions. Carbamidomethylation of Cys was used as fixed modification and oxidation of Met and deamidation of Asn and Gln as variable modifications. PeptideShaker v.1.16.15 was used to import the results for peptide and protein inference [26]. Peptide Spectrum Matches (PSMs), peptides and proteins were validated at a 1.0% False Discovery Rate (FDR) estimated using the decoy hit distribution. Only proteins having at least two unique peptides (containing at least seven amino acid residues) were considered as positively identified. The mass spectrometry proteomics data have been deposited to the ProteomeXchange Consortium via the PRIDE partner repository with the dataset identifier PXD008387 (for the extracellular vesicles data) and PXD006344 (for the ES products data).

### 2.6. RNA analyses

#### 2.6.1. mRNA and miRNA isolation

Two different biological replicates of EVs obtained from two different batches of worms were used. ODG fractions with a density between 1.07 and 1.09 (fractions containing pure EV samples after TRPS analysis) were pooled and excess Optiprep^®^ solution was removed by buffer exchanging. Total RNA and miRNA were extracted using the mirVana^TM^ miRNA Isolation Kit (ThermoFischer) according to the manufacturer’s instructions. RNA was eluted over two fractions of 50 μl each and stored at -80°C until analysed.

#### 2.6.2. RNA sequencing and transcript annotation

The RNA quality, yield, and size of total and small RNAs were analyzed using capillary electrophoresis (Agilent 2100 Bioanalyzer, Agilent Technologies, USA). Ribosomal RNA was removed from samples, which were pooled for sufficient input material for further sequencing, resulting in one sample for mRNA and two replicates for miRNA analyses, respectively. mRNA and miRNA were prepared for sequencing using Illumina TruSeq stranded mRNA-seq and Illumina TruSeq Small RNA-seq library preparation kit according to the manufacturer’s instructions, respectively. RNAseq was performed on a HiSeq 500 (Illumina, single-end 75-bp PE mid output run, approx. 30M reads per sample). Quality control, library preparation and sequencing were performed at the Ramaciotti Centre for Genomics at the University of New South Wales. The data have been deposited in NCBI's Gene Expression Omnibus and are accessible through GEO Series accession numbers GSE107985 and GSE107986.

### 2.7 Bioinformatic analyses

#### 2.7.1 Proteomics

Proteins were classified according to Gene Ontology (GO) categories using the software Blast2GO basic version 4.0.7. [27] and Pfam using HMMER v3.1b1 [28]. Putative signal peptides and transmembrane domain(s) were predicted using the programs CD-Search tool [29] and SignalP [30].

#### 2.7.2. mRNA analysis

High-throughput RNA-seq data was aligned to the *T. muris* reference genome models (WormBase WS255; http://parasite.wormbase.org; [31]) using the STAR transcriptome aligner [32]. Prior to downstream analysis, rRNA-like sequences were removed from the metatranscriptomic dataset using riboPicker-0.4.3 (http://ribopicker.sourceforge.net; [33]). BLASTn algorithm [34] was used to compare the non-redundant mRNA dataset for *T. muris* EVs to the nucleotide sequence collection (nt) from NCBI (www.ncbi.nlm.nih.gov) to identify putative homologues in a range of other organisms (cut-off: <1E-03). Corresponding hits homologous to the murine host, with a transcriptional alignment coverage <95% (based on the effective transcript length divided by length of the gene), and with an expression level <10 fragments per kilobase of exon model per million mapped reads (FPKM) normalized by the length of the gene, were removed from the list. The final list of mRNA transcripts from *T. muris* exosomes was assigned to protein families (Pfam) and GO categories (Blast2GO).

#### 2.7.3. miRNA analysis and target prediction

The miRDeep2 package [35] was used to identify known and putative novel miRNAs present in both miRNA samples. As there are no *T. muris* miRNAs available in miRBase release 21 [36], the miRNAs from the nematodes *Ascaris suum, Brugia malayi, Caenorhabditis elegans, Caenorhabditis brenneri, Caenorhabditis briggsae, Caenorhabditis remanei, Haemonchus contortus, Pristionchus pacificus, Panagrellus redivivus,* and *Strongyloides ratti* were utilised as a training set for the algorithm. Only miRNA sequences commonly identified in both replicates were included for further analyses. The interaction between miRNA and murine host genes was predicted using the miRanda algorithm 3.3a [37]. Input 3'UTR from the *M. musculus* GRCm38.p4 assembly was retrieved from the Ensembl database release 86 [38]. The software was run with strict 5' seed pairing, energy threshold of -20 kcal/mol and default settings for gap open and gap extend penalties. Interacting hits were filtered by conservative cut-off values for pairing score (>155) and matches (>80%). The resulting gene list was classified by the Panther classification system (http://pantherdb.org/) using pathway classification [39] and curated by the reactome pathway database (www.reactome.org) [40].

## 3. Results

### 3.1 Proteomics analysis of the ES products of T. muris

The ES products secreted by two different batches of *T. muris* adult worms were analysed using LC-MS/MS. A total of 1,777 and 2,056 peptide-spectrum matches (PSMs) were confidently identified in the first and second biological replicates analysed respectively. Similarly, a total of 591 and 704, corresponding to 197 and 233 proteins were identified with 100% confidence. After removing the proteins identified from only one peptide and the sequences belonging to the contaminants, 100 and 116 *T. muris* proteins were identified in both replicates. A total of 68 proteins were found in both replicates, whereas 32 and 48 proteins were uniquely found in replicate 1 and 2 respectively, resulting in 148 proteins in total (Supplementary Table 1).

The identified proteins were subjected to a Pfam and GO analysis. The most represented domains were “trypsin-like peptidase”, “thioredoxin-like” and “tetratricopeptide repeat domains”, with 21, 19 and 13 proteins containing these domains respectively (Figure 1A). The most abundant GO terms within the “molecular function” ontology were “protein binding”, “metal ion binding” and “nucleic acid” as well as “isomerase activity”, “oxidoreductase activity” and “serine-type peptidase activity” (Figure 1B). The GO terms within “biological process” and “cell component” (as well as the above mentioned “molecular function”) are detailed in Supplementary Table 1.

**Figure 1.**
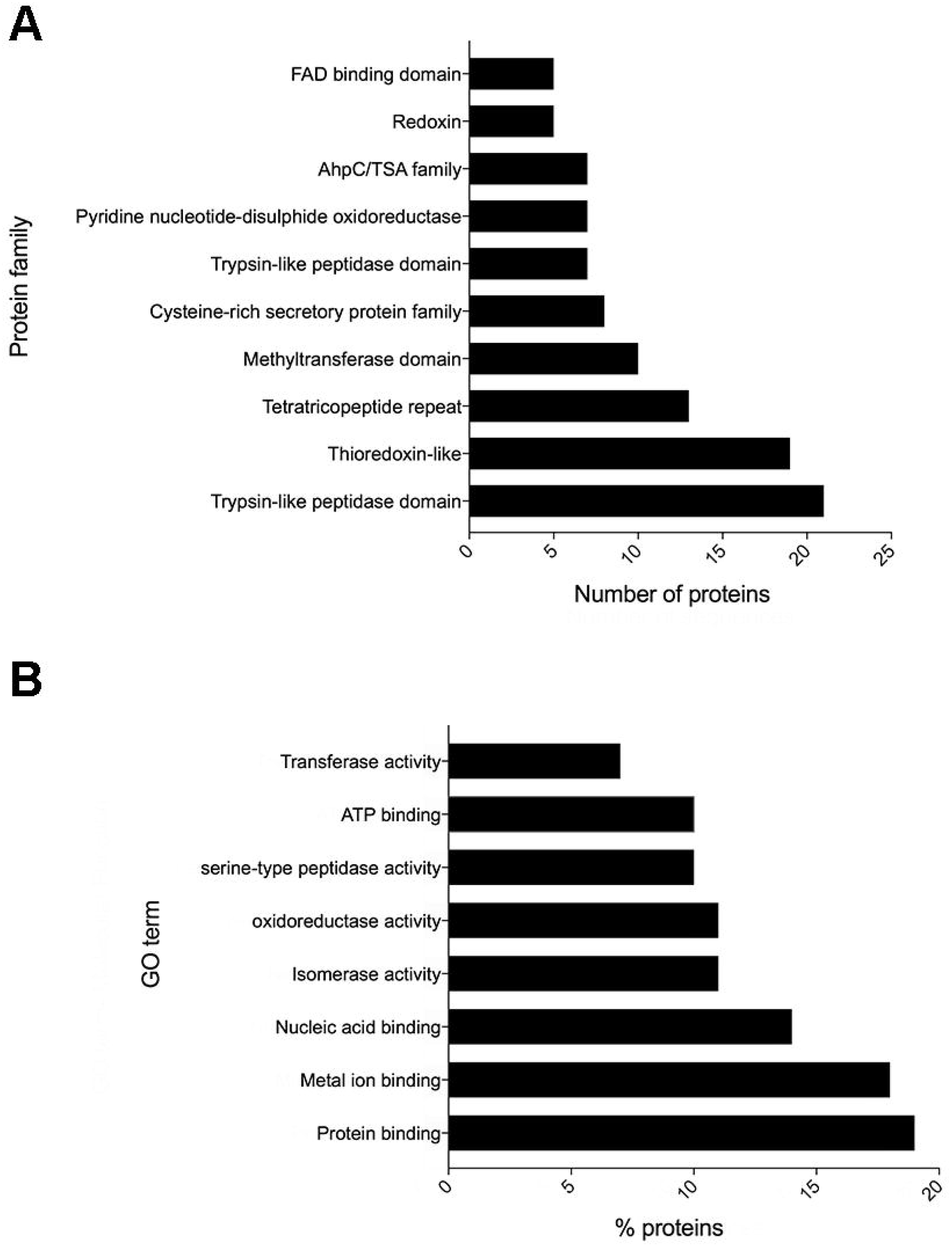
Bioinformatic analyses of the proteins secreted by *Trichuris muris*. (A) Bar graph showing the most abundant protein families after a Pfam analysis on the excretory/secretory proteins derived from *T. muris.* (B) Bar graph showing the most abundantly represented gene ontology molecular function terms in excretory/secretory proteins derived from *T. muris.*

From the total of 148 proteins found in both replicates, only 62 had a signal peptide (Supplementary Table 2), which opened the possibility of other non-classical mechanisms of secretion of these proteins described in other helminths, such as EVs.

### 3.2 T. muris adult worms secrete exosome-like EVs that can be internalised by host cells

The ES products secreted by *T. muris* adult worms were concentrated and EVs purified using Optiprep^®^ gradient. The density of the 12 fractions recovered after Optiprep^®^ separation was measured, ranging from 1.04-1.27 g/ml (Table 1). All fractions were subjected to TRPS analysis using a qNano system, but only fractions 4-10 contained enough vesicles for the analysis (Figure 2). Fraction 6 (corresponding to a density of 1.07 g/ml) contained the highest number of EVs (1.34 × 10^12^ particles/ml), followed by fractions 7 (density = 1.008 g/ml; concentration = 8.21 × 10^10^ particles/ml) and fraction 5 (density = 1.07 g/ml; concentration = 7.47 ×10^10^) (Table 1). Protein concentration was measured in all fractions, and EV purity determined as described previously [41] (Table 1). Fraction 6 had the purest EV preparation (4.31× 10^9^ particles/μg), followed by Fractions 7 and 5 (4.04 ×10^8^ and 1.58 ×10^8^ particles/μg respectively) (Table 1). Furthermore, the vesicle size was determined using the qNano system, and the results are summarised in Table 1.

**Table 1.** Features of the different fractions isolated after Optiprep fractionation of extracellular vesicles from *Trichuris muris.* Despite protein being detected in all fractions, only vesicles from fractions 4-10 could be quantified. The purity of the different fractions was calculated according to [41].

| **Optiprep fraction** | **Density (g/mL)** | **Protein quantification (µg/ml)** | **Particle concentration (particles/mL)** | **Purity of vesicles particles/µg** | **Particle size nm** |
| --- | --- | --- | --- | --- | --- |
| **S1** | 1.04 | 104.69 | - | - | - |
| **S2** | 1.05 | 290.10 | - | - | - |
| **S3** | 1.05 | 435.58 | - | - | - |
| **S4** | 1.06 | 477.86 | 2.96E+08 | 6.19E+05 | 93±41.5 |
| **S5** | 1.07 | 474.19 | 7.47E+10 | 1.58E+08 | 72±11.7 |
| **S6** | 1.07 | 310.61 | 1.34E+12 | 4.31E+09 | 72±23.8 |
| **S7** | 1.08 | 202.99 | 8.21E+10 | 4.04E+08 | 90±25.5 |
| **S8** | 1.10 | 160.76 | 1.31E+10 | 8.15E+07 | 152±75.3 |
| **S9** | 1.12 | 190.46 | 1.31E+10 | 6.88E+07 | 183±68.2 |
| **S10** | 1.15 | 436.86 | 1.71E+09 | 3.91E+06 | 165±54.9 |
| **S11** | 1.21 | 232.77 | - | - | - |
| **S12** | 1.27 | 83.35 | - | - | - |

**Figure 2.**
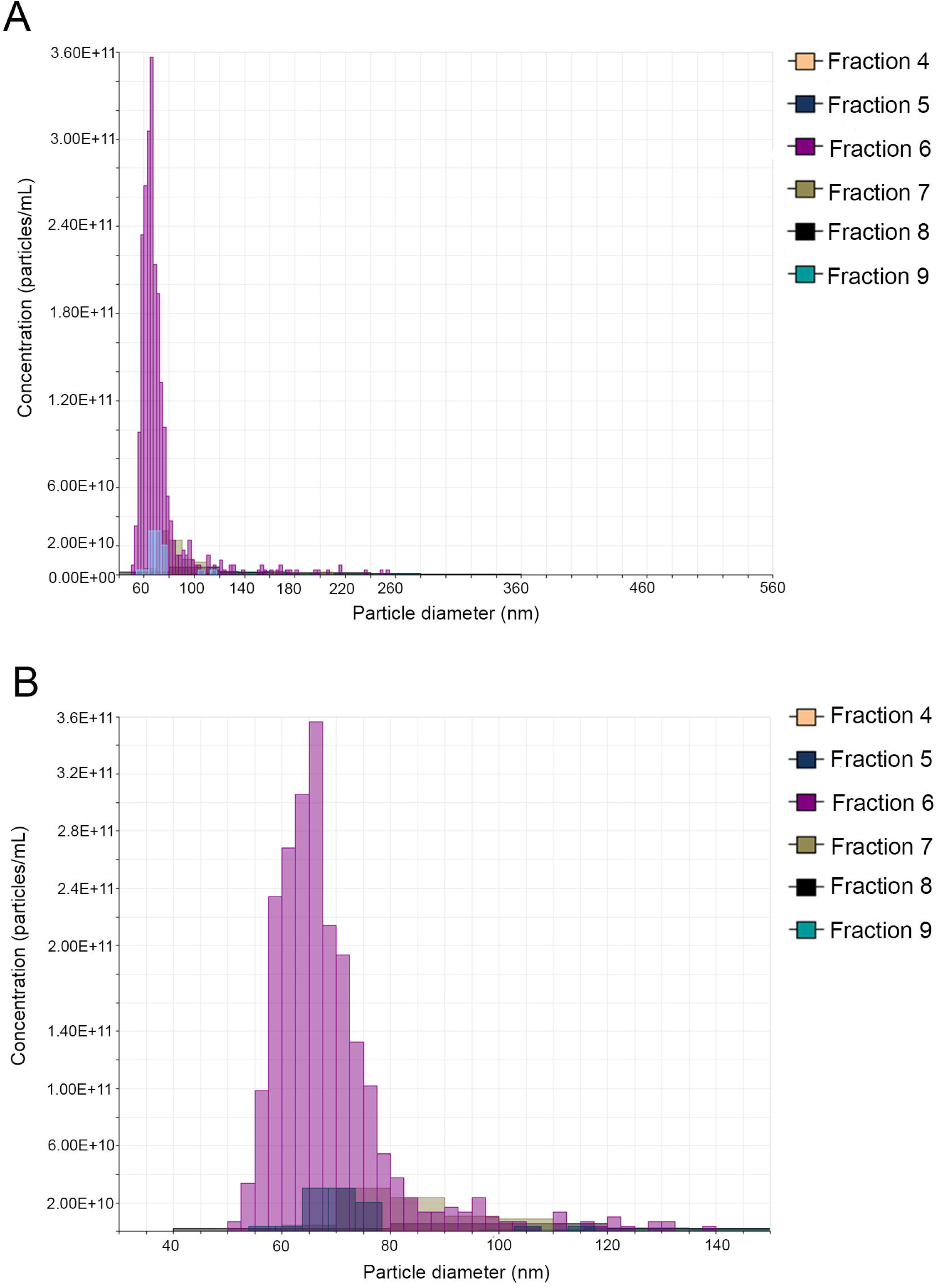
Tunable resistive pulse sensing analysis of the extracellular vesicles (EVs) secreted by *Trichuris muris*. (A) The size and number of the EVs secreted by *T. muris* after purification using an Optiprep^®^ gradient was analysed using a qNano system (iZon). (B) Detailed graph showing number of vesicles with a diameter between 30-150nm. Only fractions 4-10 contained enough vesicles for the analyses.

In vitro cellular uptake of *T. muris* EVs in host cells was studied using murine colonic organoids, comprised of the complete census of progenitors and differentiated cells from the colon epithelial tissue growing in cell culture. Purified membrane-labelled *T. muris* EVs were injected into the central lumen of the colonic organoids (corresponding to the intestinal lumen), and they were incubated for 3 hours at 37°C to demonstrate cellular uptake. We observed internalisation of EVs by organoid cells, which was absent by preventing endocytosis in metabolic inactive cells at 4°C (Figure 3). Using confocal microscopy, we confirmed that EVs were inside cells and present a cytoplasmic location of the stained EVs in some cells within the donut-shaped epithelial layer. By analysing the pictures and comparing the area of epithelial cells and the central lumen uptake of stained vesicles at 37°C could be quantitatively traced within epithelial organoid cells (mean percentage of corrected total fluorescence (%CTF) +/-SD: 4.04 +/-1.11), and %CTF values were significantly reduced (p<0.001) in the central lumen (0.59 +/- 0.41), whereas at 4°C, CTF values were 0.21 +/- 0.29 and 3.79 +/-2.29 for the total epithelial organoid cells and the central lumen, respectively (Figure 3E).

**Figure 3.**
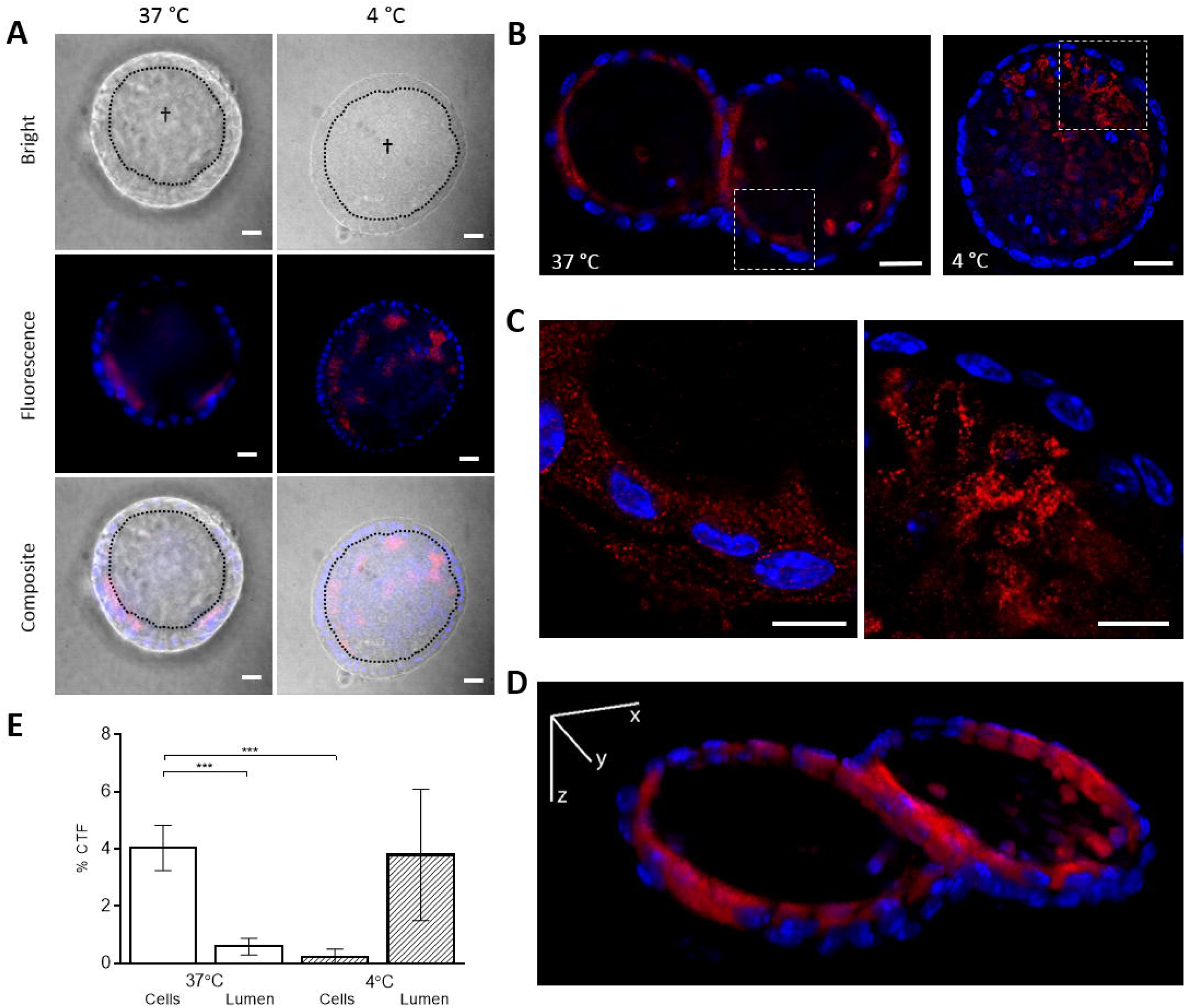
*Trichuris muris* extracellular vesicles (EVs) are internalized by murine colonic organoids. (A) Representative fluorescence images (Zeiss AxioImager M1 ApoTome) of PKH26-labelled EVs (red) internalized by organoids after 3 hours at 37°C and 4°C (metabolically inactive cells). Hoechst dye (blue) was used to label cell nuclei. † indicates the lumen of the organoids, which corresponds to the murine gut lumen, separated by the dotted line from the epithelial cell layer. White bar corresponds to 20 μm. (B) Deconvolved laser scanning confocal microscopy images (Zeiss 780 NLO) of murine colonic organoids under 20x magnification. White bar corresponds to 20 μm. (C) Magnification of the framed area in B under 100x magnification. White bar corresponds to 10 μm. (D) 3D projection of z-stack serial confocal images of a 12 μm organoid slice incubated with PKH26-labelled EVs after 3 hours at 37°C (videos of 3D projections of the experiments at 37°C and 4°C are available in the supplementary materials) (E) Percentage of the corrected total fluorescence (CTF) adjusted by background fluorescence and the surveyed area of PKH26-stained EVs in total epithelial cells (donut-shaped selection) or in the organoid lumen incubated under different conditions in 10 different organoids from 2 technical replicates (5 each). *** indicates highly significant results (p<0.001). Error bars indicate 95% confidence intervals.

### 3.3 T. muris secreted EVs contain specific proteins

Two replicates containing ODG fractions with a density between 1.07-1.09 g/ml were subjected to SDS-PAGE separation, each lane cut into 9 slices and subjected to trypsin digestion followed by LC-MS/MS analysis. The results obtained from both replicates were combined and only proteins commonly found in both replicates were considered for further analysis. A total of 28,376 and 19,510 spectra corresponding to 6,489 and 5,455 peptides were identified in replicates 1 and 2 respectively. A total of 663 and 718 proteins matching *T. muris, M. musculus* and common contaminants for the cRAP database were identified (Supplementary Table 3). From these, 465 and 26 proteins containing at least 2 validated unique peptides matched *T. muris* and *M. musculus respectively in replicate 1. Similarly,* 486 and 36 proteins containing at least 2 validated unique peptides matched *T. muris* and *M. musculus respectively in replicate 2.* A final list of 364 and 17 proteins corresponding to *T. muris* and *M. musculus* respectively was defined with proteins commonly identified in both replicates (Supplementary Table 4). Only these common proteins were used in subsequent analysis

Among the identified proteins from *T. muris,* the most abundant proteins based on the spectrum count were a trypsin domain-containing protein, several sperm-coating protein (SCP)-like extracellular proteins, also called SCP/Tpx-1/Ag5/PR-1/Sc7 domain containing proteins (SCP/TAPS), a poly-cysteine and histidine-tailed protein, a glyceraldehyde-3-phospahte dehydrogenase and a TB2/DP1 HVA22 domain-containing protein. One tetraspanin (TMUE_s0037005100) was found in the EVs sample, as well as other proteins typically found in EVs from helminths like 143-3, heat shock protein (HSP) and glutathione-s-transferase were also identified in this study. Furthermore, from the 364 identified proteins from *T. muris,* only 50 (13.7%) contained a transmembrane domain, and 120 (32.96%) had a signal peptide. Despite washing the worms extensively before culturing, discarding the first 4 hours of the ES for EV isolation (which typically contains a significant amount of host proteins) and analysing only fractions containing highly pure EV samples, we found some host proteins in our analysis. Among these proteins we found an antibody Fc fraction and 9 proteasome proteins (Supplementary Table 4).

Following a GO analysis, the most represented GO terms within biological process in the *T. muris* EV proteins were assigned as “cellular metabolic process”, “response to stimulus” and “proteolysis” (Figure 4A). Similarly, the most represented GO terms within molecular function were “Oxidoreductase activity”, “protein binding” and “ATP binding” (Figure 4B).

**Figure 4.**
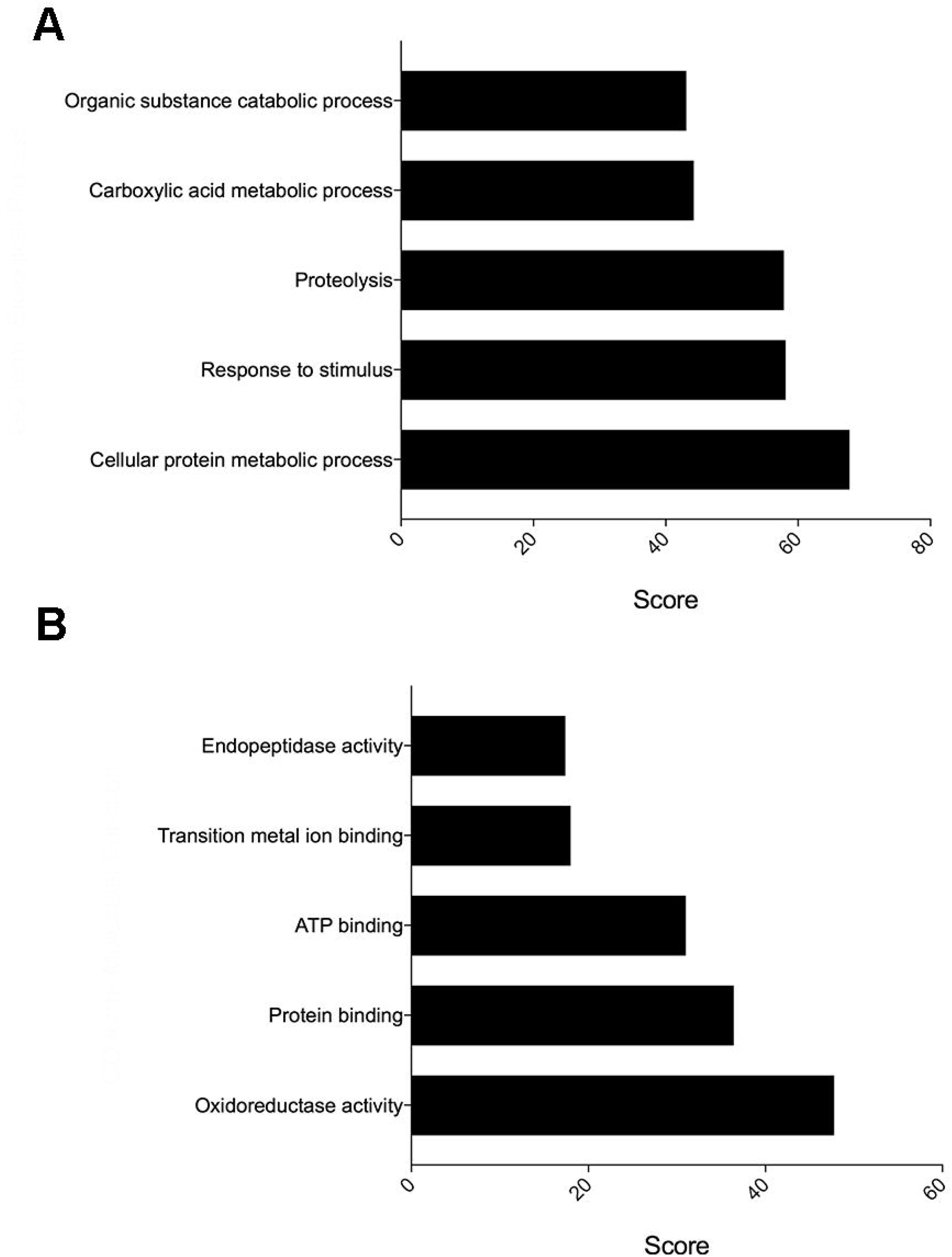
Gene ontology analysis of proteins from the extracellular vesicles (EVs) secreted by *Trichuris muris*. (A) Bar graph showing the most abundantly represented gene ontology biological process terms in proteins present in the EVs secreted by *T. muris.* (B) Bar graph showing the most abundantly represented gene ontology molecular function terms in proteins present in the EVs secreted by *T. muris.*

### 3.4 T. muris secreted EVs contain specific mRNAs and miRNAs

RNA content of EVs was characterized using the Illumina HiSeq platform. For an initial description of parasite-specific mRNAs in EVs, total RNA from a highly pure EV sample was sequenced and results were curated based on stringent thresholds. This resulted in 475 full-length mRNA transcripts mapping to *T. muris* gene models. The identified hits were subjected to a Pfam and GO analysis. Interestingly, the most represented domains were of “unknown function”, “reverse transcriptase”, and “helicase”, whereas other gene models with DNA-binding and processing domains were also highly abundant (e.g. genes with “retrotransposon peptidase” domain) (Figure 5 A). Mapping to molecular functions identified “protein binding” as the most abundant term, with 31.9% of all sequences involved in this function (Figure 5B). The underlying proteins from the parasite-specific mRNAs had functions in signalling and signal transduction, transport, protein modification and biosynthetic processes, as well as in RNA processing and DNA integration (Figure 5C). Data is provided in Supplementary Table 5.

**Figure 5.**
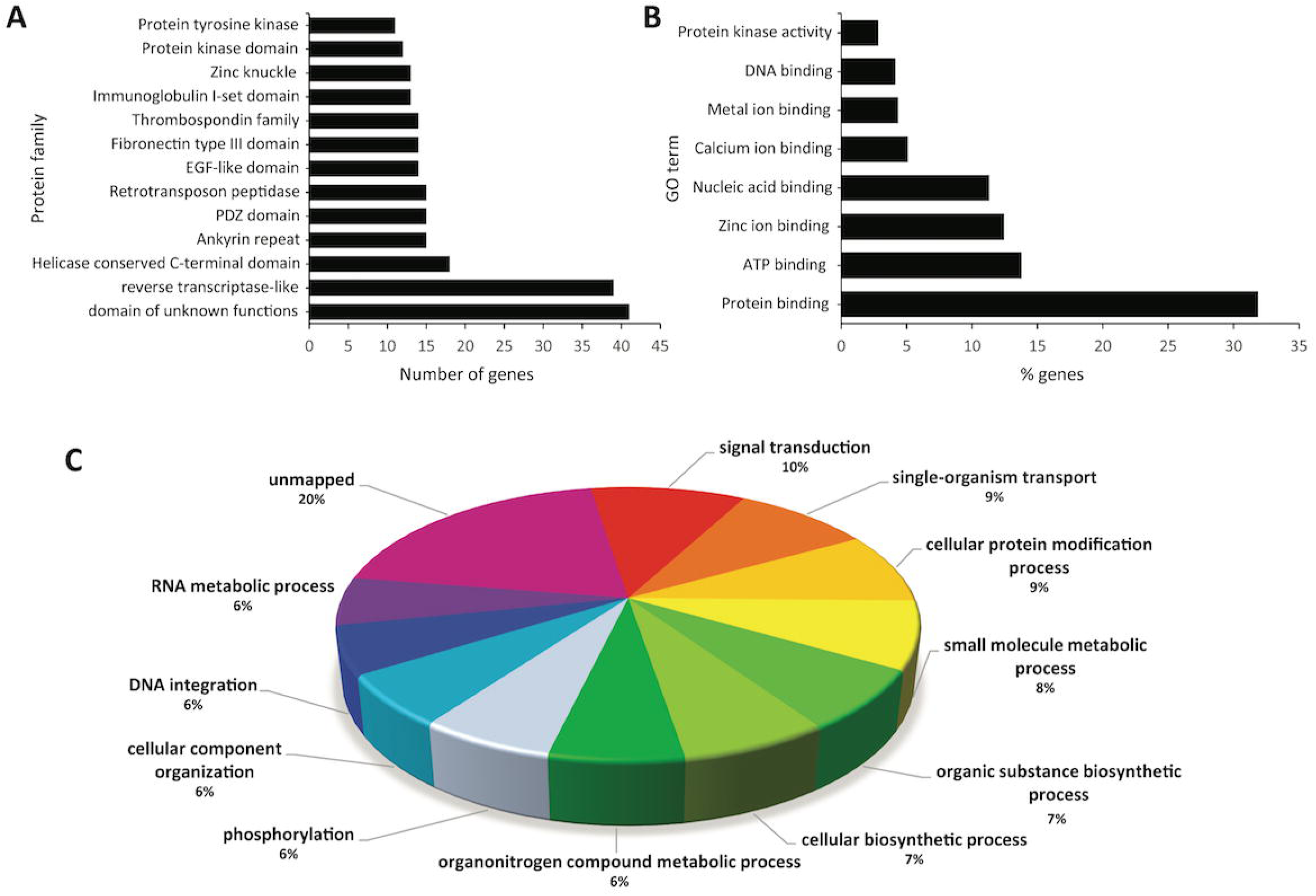
Analysis of the 475 full-length mRNAs detected in *Trichuris muris* extracellular vesicles. (A) Bar graph showing the most represented protein families (Pfam) from the translated mRNAs. (B) Molecular functions and (C) biological functions of proteins encoded by each of the 475 transcripts assigned to gene ontology functional annotation.

By sequencing and screening biological duplicates for miRNAs, we identified 56 miRNAs commonly present in both datasets, 34 of which having close homologs in other nematodes. The remaining 22 miRNAs were novel and were named serially according to their mean abundances (tmu.miR.ev1 to tmu.miR.ev22). Potential interactions of *T. muris* miRNAs to murine host genes were explored by computational target prediction. The 56 nematode EV-miRNAs were predicted to interact with 2,043 3’UTR binding sites of the mouse genome assembly (Supplementary Table 6). Associated annotated coding genes were grouped according to signalling, metabolic, and disease pathways (Supplementary Figure 1). Indeed, a number of the nematode miRNA-mouse gene interactions are involved in host immune system, receptor, and transcriptional regulation (Figure 6). Within the 56 identified EV miRNAs, 3 (5.4%) could not be assigned for interaction with a specific pathway in the murine host, including the second most abundant asu-miR-5360-5p.

**Figure 6.**
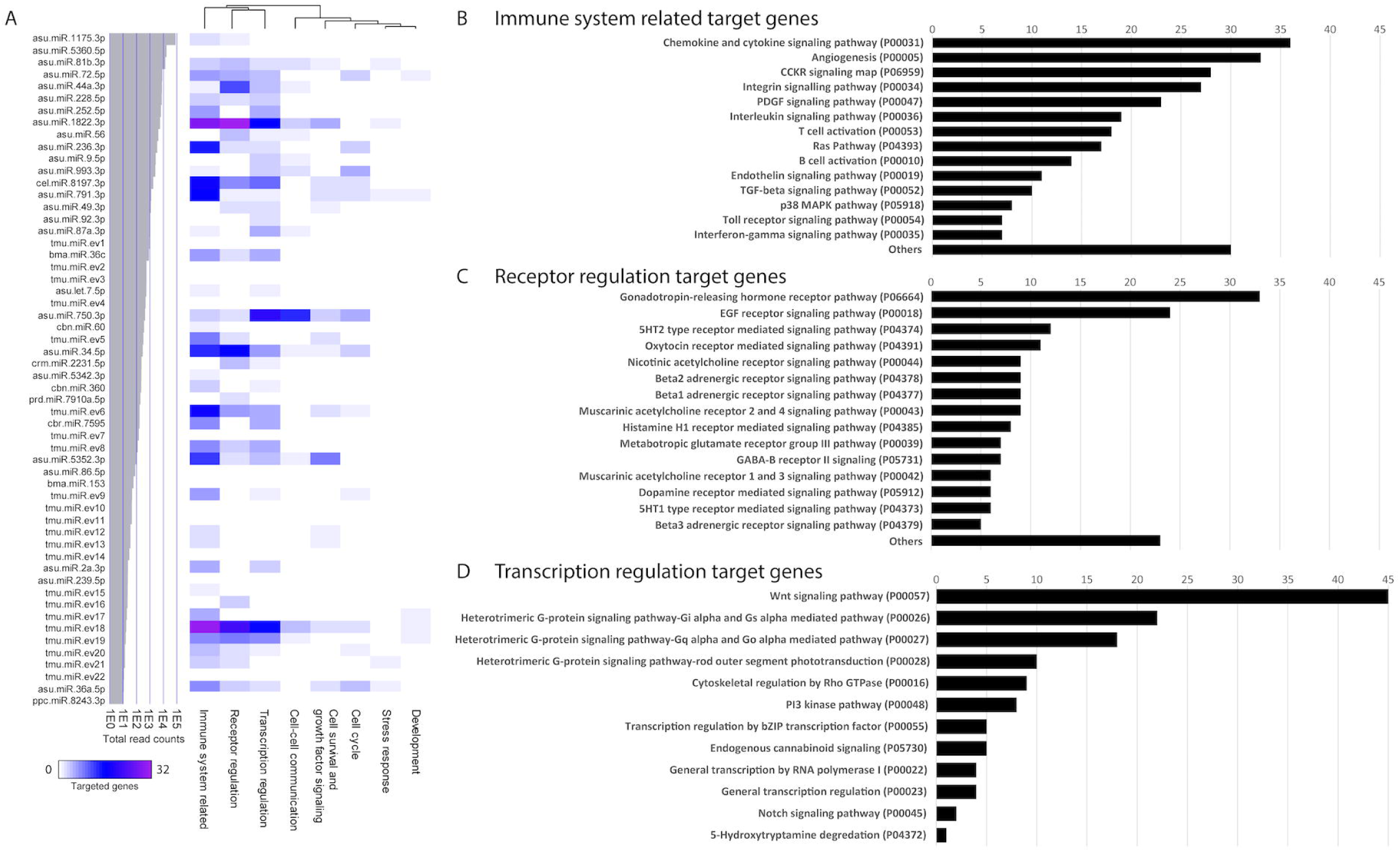
Prediction of *T. muris* extracellular vesicle (EV) miRNA target interactions to murine host genes. Functional map of *T. muris* EV miRNAs and their target murine host genes. (A) Individual targeted host genes are categorized by PantherDB signaling pathway analysis (heat map corresponds to individual targeted genes in the murine host). Bottom axis shows the 56 identified miRNAs in *T. muris* EVs and their abundances (average mean read counts from two biological replicates), termed according to their closest homologues *(de novo* transcripts were designated as tmu.miR.ev#). Total number of targeted genes identified by PantherDB categories classified as (B) ‘immune system related’, (C) ‘receptor regulation’, and (D) ‘transcription regulation’.

## 4. Discussion

Trichuriasis is a soil transmitted helminth infection that affects almost 500 million people worldwide [1, 42, 43]. In addition to the pathogenicity associated with the disease, the infection can also cause physical and intellectual retardation [1, 44]. There is, therefore, an urgent need to understand the mechanisms by which the parasite interacts with its host such that novel approaches to combat this neglected tropical disease can be developed [45]. *T. trichiura* is the main species that affects humans, but the difficulty in obtaining worms and working with the adult stage have prompted parasitologists and immunologists to use the *T. muris* rodent model.

We provide herein the first high throughput study of the secretome of *T. muris.* The analysis of the genome from *T. muris* predicted 434 proteins containing signal peptides [8]. We have confidently identified (with 2 or more peptides) 148 proteins secreted by adult *T. muris,* corresponding to 34.1% of the total predicted secreted proteins [8]). From the total proteins identified, 68 were commonly found in both replicates, highlighting the importance of analysing multiple batches of samples when conducting proteomics analyses of parasitic ES products. Among the identified proteins we found several peptidases and proteases (such as serine proteases, pepsin and trypsin domain-containing proteins) and also protease inhibitors including WAP domain-containing proteins. These findings are in agreement with the functional annotation of the *T. muris* proteins predicted from the genome [8]. Protease inhibitors (particularly serine protease inhibitors and secretory leukocyte proteinase inhibitor (SLPI)-like proteins - proteins containing mostly WAP domains) are abundantly represented in the *T. muris genome* [8]. SLPI-like proteins have been suggested to have immunomodulatory properties as well as a role in wound healing [8, 46, 47, 48], so they could be secreted in an attempt to modulate the host’s immune response and repair damage caused by both feeding/migrating worms and immunopathogenesis. In addition, we found five SCP/TAPS (also known as CAP-domain) proteins. SCP/TAPS proteins are abundantly represented in soil-transmitted helminths, although they have not been well characterised in the clade I nematodes [49].

Only recently, different authors have shown the importance of helminth-secreted EVs in host-parasite interactions. The secretion of small EVs was demonstrated in various intracellular and extracellular parasites, interacting with their hosts in a specific manner (reviewed in [17]). In addition, the secretion of EVs has been demonstrated thus far only in a small number of nematodes, including the free-living *C. elegans,* the filarial nematodes *Brugia malayi* and *Dirofilaria immitis,* the rodent nematode *Heligmosomoides polygyrus* and the ovine and porcine intestinal nematodes *Teladorsagia circumcincta* and *Trichuris suis,* respectively [20, 50, 51, 52, 53].

Our results show that *T. muris* secretes EVs with a wide variety of sizes (40-550 nm). In order to study the exosome-like vesicles (vesicles with a size between 50-150 nm) and eliminate contamination with soluble proteins that could be co-precipitated in the ultracentrifugation step, we further purified the EVs using Optiprep and analysed only fractions 5-7 (fractions containing EVs with sizes between 72±23.8nm to 90±25.5nm). For a totally novel approach in EV research, we introduced and established a long-term primary *in vitro* culture to generate 3D intestinal organoids, recapitulating the *in vivo* epithelial tissue organisation and representing the complete census of progenitors (stem cells) and differentiated cells [22, 54]. Although there are colonic cancer cell lines available, such as the intestinal epithelial cell line Caco2, cell lines cannot recapitulate the complex spatial organisation of the intestinal epithelium, they have undergone significant molecular changes to become immortal, and do not represent all intestinal subsets [55]. Hence, we used colonic organoids corresponding to the epithelial barrier, which is the first line of defence against intestinal pathogens. In a first attempt to study whipworm EV interactions with the host intestinal epithelial barrier, we observed vesicle uptake only in a subset of cells. This could not be confirmed by repeating the experiment and analysing the cells by laser scanning confocal imaging and z-stack rendering. At first glance, this suggests that parasite EV uptake - at least under the tested conditions - is a cell type-unspecific process. Results from several studies show that fluorescently labelled EVs can be taken up by different cell types, whereas other studies indicate that vesicular uptake is a highly cell-type specific process (summarized in [56]). As we don’t know which intestinal cells are presented in the screened murine colonic organoids, further studies should include good host cell markers to distinguish the different cells in the heterogeneous organoid system. Furthermore, as the tested conditions could correspond to a high parasite burden present in severe infections, titration of the administered EV dosages, and inclusion of different endocytosis-inhibitors could give clues about the specificity and mechanisms of *T. muris* EV-host interactions. In line with the observation of different studies, uptake of EVs was dramatically reduced when incubating at 4°C, suggesting that internalization is not a passive process occurring in metabolically inactive cells, and hence relies on some source of energy [57, 58, 59, 60]. A disadvantage of the intestinal organoid culture is the lack of any immune cells. Co-culture experiments with intestinal organoids and intraepithelial lymphocytes as described by Nozaki and colleagues [61] could be a powerful tool to study interactions of EVs with immune cells at their primary interface, complemented with host cell proteomics to detect host and parasite proteins, and transcriptomic studies.

The proteomic analysis of the exosome-like EVs showed a total of 381 proteins (364 from *T. muris* and 17 from the host), 130 of which have been also found in the crude ES prep. From the common proteins, only 54 (41.5%) were predicted to have a signal peptide, thus, EVs could be a potential mechanism by which these proteins are secreted by helminths into the extracellular milieu, addressing an issue that has been frequently debated in the literature [8]. It is interesting to note that one tetraspanin (TMUE_s0037005100) was detected in the *T. muris* EVs. Tetraspanins are considered a molecular marker of exosomes since they are present on the surface membrane of EVs from many different organisms including mammalian cells and bacteria [62]. EVs secreted by, or shed from the surface of parasitic trematodes are enriched in tetraspanins [18, 21, 63], although, in the case of nematodes, they are not abundant on the surface of EVs. For instance, only one member of this family was found in the EVs secreted by the nematode *H. polygyrus* [20]. Since tetraspanins are involved in the formation of the membrane of EVs [62], it is unclear why EVs secreted by nematode parasites are not replete in tetraspanins, as happens in trematodes. In trematodes, exosomes derive from the tegumental syncytium of the worm [64], whereas in nematodes they seem to have an intestinal origin [20]. This different origin could be the reason why tetraspanins are not enriched in nematode EVs. Our dataset presents other proteins usually found in parasitic exosomes, such as 14-3-3, heat shock protein 90 and myoglobin.

Proteins involved in proteolysis were abundantly represented (12.3% of sequences) in the *T. muris* EVs (e.g. trypsin like, cathepsins and aminopeptidases). *Trichuris* lacks the muscular pharynx that many other nematodes use to ingest their food, a challenging process given the hydrostatic pressure of the pseudocoelom that characterizes the phylum. Instead, it has been suggested that the parasite secretes copious quantities of digestive enzymes for this purpose [8]. We have shown that proteases are heavily represented in the ES products, and proteolysis is also in the top 3 main GO terms found when we analysed the proteins present in the EVs. Indeed, 37 of the 364 proteins from *T. muris* found in the EVs contain a trypsin or trypsin-like domain. These proteins could be involved in extracellular digestion, and, since feeding is a key process in parasite biology, they might also be potential targets for vaccines and drugs against the parasite. Helminth proteases have also been hypothesized to be involved in immunomodulatory processes, where they degrade important immune cell surface receptors [65] and host intestinal mucins [9, 66]. If this is the case, *Trichuris* could be secreting EVs containing peptidases to promote an optimal environment for attaching to the mucosa and feeding purposes.

Proteins containing an SCP/TAPS domain were identified in the EVs secreted by *T. muris*. This family of proteins is abundantly expressed by parasitic nematodes and trematodes. For instance, they represent 35% of the ES products of the hookworm *Ancylostoma caninum [67],* and have been found in free-living and plant nematodes (reviewed by [68]). Their role is still unknown, although they have been suggested to play roles in fundamental biological processes such as larval penetration [69], modulation of the immune response [70, 71], in the transition from the free-living to the parasitic stage [72] and have even been explored as vaccine candidates against hookworm infections [73]. It is interesting to note that EVs from other helminths are enriched for many known vaccine candidate antigens [21]. Since SCP/TAPS proteins are abundant in the EVs secreted by *T. muris,* their potential use as vaccines should be further explored.

We analysed the mRNA and miRNA content of the exosome-like EVs secreted by *T. muris* since it has been well documented that the nucleic acid content of eukaryotic EVs can be delivered between species to other cells, and can be functional in the new location [74]. Functional categorization of the 475 mRNAs from *T. muris* EVs revealed a high proportion of protein-binding proteins. Interestingly, mRNAs for common EV proteins were present, including inter alia mRNAs for tetraspanins, HSPs, histones, ubiquitin-related proteins, and signalling- and vesicle trafficking molecules (rab, rho and ras). A significant number of domains found in the proteins predicted from mRNA sequences were involved in reverse transcription and retrotransposon activity, suggesting a strong involvement of these mRNAs in direct interactions with the host target cell genome. This is supported by the hypothesis of shared pathways between EV biogenesis and retrovirus budding, including the molecular composition of the released particles, sites of budding in different cell types, and the targeting signals that deliver proteins [75, 76, 77, 78].

To gain a more comprehensive picture of the RNA composition of the *T. muris* EVs, we sequenced the miRNAs present in *T. muris* EVs and identified 56 miRNAs, including 22 novel miRNAs without described homology to other nematodes. We also identified miRNAs that shared homology with those from other parasitic nematodes, such as let-7, miR-2, miR-9, miR-34, miR-36 (a and c), miR-44, miR-60, miR-72, miR-81, miR-86, miR-87, miR-92, miR-228, miR-236 and miR-252 (reviewed in [79]). This suggests that secretion of miRNAs by parasitic nematodes is most probably conserved and that EVs could be playing an important role in this secretory pathway. *T. muris* miRNAs that regulate expression of genes involved in specific conditions and cellular pathways were identified. In humans, more than 60% of all protein-coding genes are thought to be controlled by miRNAs (reviewed in [80]). Our *in silico* prediction analysis of murine host gene interactions of *T. muris* EV miRNAs points towards a strong involvement of parasite miRNAs in regulation/modulation of the host immune system [81]. In this sense, it has been previously demonstrated that small EVs secreted by *H. polygyrus* interact with intestinal epithelial cells of its murine host and suppress type 2 innate immune responses, promoting parasite survival [20]. Similarly, other studies demonstrated the secretion of EVs containing miRNAs by larvae of the porcine whipworm *T. suis*, and although the miRNAs were not sequenced, the authors suggested a possible role in immune evasion [50].

The mechanisms by which parasitic helminths pack their nucleic acid cargo into EVs is still unknown, and, while we hypothesize that an active mechanism might regulate this process, we cannot discard the possibility that mRNAs and miRNAs could be internalised at random. Understanding this mechanism will be of importance in understanding the intimately interactive nature of host-parasite biology. For example, are mRNAs in parasite EVs translated into protein by target host cells, akin to viral hijacking of host cell protein manufacturing machinery? Or, are EV mRNAs unimportant, and manipulation of host cell gene expression is mostly due to miRNAs?

In the present study we have provided important information regarding the molecules secreted by the murine whipworm *T. muris.* The identification of the secreted proteins and EVs (including their proteomic and RNA content) will prove useful not only for the design of novel approaches aimed at controlling whipworm infections, but also to understand the way the parasite promotes an optimal environment for its survival.

---

Supplementary Figure 1. Prediction of *T. muris* extracellular vesicle (EV) miRNA target interactions to murine host genes. Functional map of *T. muris* EV miRNAs and their target murine host genes categorized by PantherDB signalling, metabolic, disease, and other pathways. Heat map corresponds to individual targeted genes in the murine host.

Supplementary Table 1. Detailed analysis of the proteins secreted by *Trichuris muris.* Proteins were annotated using Blast2GO [27], and the description blast e-values, gene ontology (GO) terms and enzyme codes are shown. Information about the number of unique peptides, peptide-spectrum matches (PSM), spectrum counting and coverage is also provided for each protein and batch analysed.

Supplementary Table 2. Signal peptide analysis of the proteins secreted by *Trichuris muris.* An analysis to check for the presence of a signal peptide in each of the proteins secreted by *T. muris* was performed using SignalP [30].

Supplementary Table 3. Proteomic analysis of extracellular vesicles (EVs) secreted by *Trichuris muris.* Details of the identification of the proteins present in the EVs secreted by *T. muris* using X!Tandem, Tide, MS-GF + and OMSSA. All proteins are shown, including contaminants, and independently of the number of unique peptides identified.

Supplementary Table 4. Curated proteomic analysis of extracellular vesicles (EVs) secreted by *Trichuris muris.* Proteins from *T. muris* and *Mus musculus* found in both replicates of EVs secreted by *T. muris.* Only proteins found in both replicates containing at least 2 validated peptides were considered for the analysis. Proteins are sorted by theoretical abundance (average spectrum counting). Only proteins from *T. muris* or *M. musculus* are shown.

Supplementary Table 5. Data on 475 detected mRNA transcripts from *T. muris* extracellular vesicles. RNA-seq data was aligned to the *T. muris* reference genome models [31] and the E-value of the alignment, the fragments per kilobase of exon model per million mapped reads (FPKM) normalized by the length of the gene and the relative coverage of the alignment are provided.

Supplementary Table 6. Data description on predicted *T. muris* miRNA-host target interactions. Table showing the 56 miRNAs identified in the *T. muris* extracellular vesicles and their 2,043 3’UTR predicted binding sites in the mouse genome.

Supplementary Video 1. 3D reconstruction of a 12 μm murine colonic organoid slice incubated with PKH26-labelled EVs (red) after 3 hours at 37°C. Z-stacks serial images acquired by laser scanning confocal microscopy under 20x magnification and processed in ImageJ software. Cell nuclei are stained with Hoechst dye (blue).

Supplementary Video 2. 3D reconstruction of a 10 μm murine colonic organoid slice incubated with PKH26-labelled EVs (red) after 3 hours at 4°C. Z-stacks serial images acquired by laser scanning confocal microscopy under 40x magnification and processed in ImageJ software. Cell nuclei are stained with Hoechst dye (blue).

